# Ancient DNA Insights into Aboriginal Australian Mortuary Practices

**DOI:** 10.1101/2020.06.11.145482

**Authors:** Sally Wasef, Joanne L Wright, Shaun Adams, Michael C Westaway, Clarence Flinders, Eske Willerslev, David Lambert

**Affiliations:** Australian Research Centre for Human Evolution, Environmental Futures Research Institute, Griffith University, 170 Kessels Road, Nathan 4111, Australia; Archaeology, School of Social Science, University of Queensland, Sir Fred Schonell Drive, St Lucia QLD 4072; Cape Melville, Flinders and Howick Islands Aboriginal Corporation, Cairns, Australia; Department of Zoology, University of Cambridge, Cambridge, UK; Wellcome Trust Sanger Institute, Cambridge, UK; Lundbeck Foundation GeoGenetics Center, University of Copenhagen, Copenhagen, Denmark

**Keywords:** Aboriginal Australians, bioarchaeology, genomic enrichment, mitochondrial DNA, paleogenetics, Ancient DNA

## Abstract

Paleogenetics is a relatively new and promising field that has the potential to provide new information about past Indigenous social systems, including insights into the complexity of burial practices. We present results of the first ancient DNA (aDNA) investigation into traditional mortuary practices among Australian Aboriginal people with a focus on North-East Australia. We recovered mitochondrial and Y chromosome sequences from five ancestral Aboriginal Australian remains that were excavated from the Flinders Island group in Cape York, Queensland. Two of these individuals were sampled from disturbed beach burials, while the other three were from bundle burials located in rock shelters. Genomic analyses showed that individuals from all three rock shelter burials and one of the two beach burials had a close genealogical relationship to contemporary individuals from communities from Cape York. In contrast the remaining male individual, found buried on the beach, had a mitochondrial DNA sequence that suggested that he was not from this location but that he was closely related to people from central Queensland or New South Wales. In addition, this individual was associated with a distinctive burial practice to the other four people. It has been suggested that traditionally non-locals or lower status individuals were buried on beaches. Our findings suggest that theories put forward about beach burials being non-local, or less esteemed members of the community, can potentially be resolved through analyses of uniparental genomic data. Generally, these results support the suggestion often derived from ethnohistoric accounts that inequality in Indigenous Australian mortuary practices might be based on the status, sex, and/or age of individuals and may instead relate to place of geographic origin. There is, however, some departure from the traditional ethnohistoric account in that complex mortuary internments were also offered to female individuals of the community, with genomic analyses helping to confirm that the gender of one of the rockshelter internments was that of a young female.

## 1 Introduction

A better understanding of how people lived in the past can be revealed by an examination of their skeletal remains, which can assist in reconstructing their life history. The examination of their burial context reveals information on how they were treated after death. Traditionally this level of understanding has been reconstructed using methods from biological anthropology. These methods can include studies of craniometrics and biological traits that can provide insights into the genetic relatedness of an individual Pardoe (1993). Moreover, the examination of the health status of an individual which provides insights into how the individual may have been nursed or cared for by that society during life (Tilley, 2015). Taphonomic investigations of burial sites and the arrangement of the deceased in the grave provide important information on the status of an individual (Pearson, 1999). However, the methods based on archaeological and anthropological assessments are sometimes insufficient to establish the sex and biological kinship relationships particularly when ancestral remains are heavily eroded or damaged and when comparative datasets simply do not exist for multivariate analyses of metric and non-metric data. The uses of ancient DNA (aDNA) can provide more precise information about the biological affinity among individuals in past populations to complement the other bioarcheological findings. The field of ancient DNA has expanded rapidly since its inception in 1984 (Kutanan et al., 2017). It has now impacted a large number of different disciplines including biological anthropology. Ancient DNA studies have the potential to provide new information about cultural traditions and specifically burial practices.

Biparentally inherited genomic data has been successively used in population studies for repatriation purposes (Heupink et al., 2016; Malaspinas et al., 2016). However, due to the highly fragmented nature of aDNA, this whole-genome approach could be inefficient when studying the burial practices of poorly preserved remains (Collard et al., 2019). It is typically more feasible to recover high copy number uniparental mitochondrial DNA (mtDNA) from human remains (Wright et al., 2018).

The Flinders Island group is located in the tropical north-east of Queensland, off the eastern coast of Cape York Peninsula. The group consists of seven continental Islands, sitting within the Princess Charlotte Bay, west from Cape Melville (**Figure 1**). Over the past two centuries, Indigenous people from these Islands have been regularly involved in maritime industries. Cultural changes that have included increased European contact have also resulted in the removal and theft of many Indigenous cave bundle burials (Horsfall, 1991). Not only were ancient remains affected by these activities, but so too were the Indigenous people of Flinders themselves. Since the Second World War many Indigenous people were forcefully removed from Flinders Island (Wurrima) to the mainland as a result of government regulations at that time (Rigsby & Chase, 1998).

**Figure 1.**
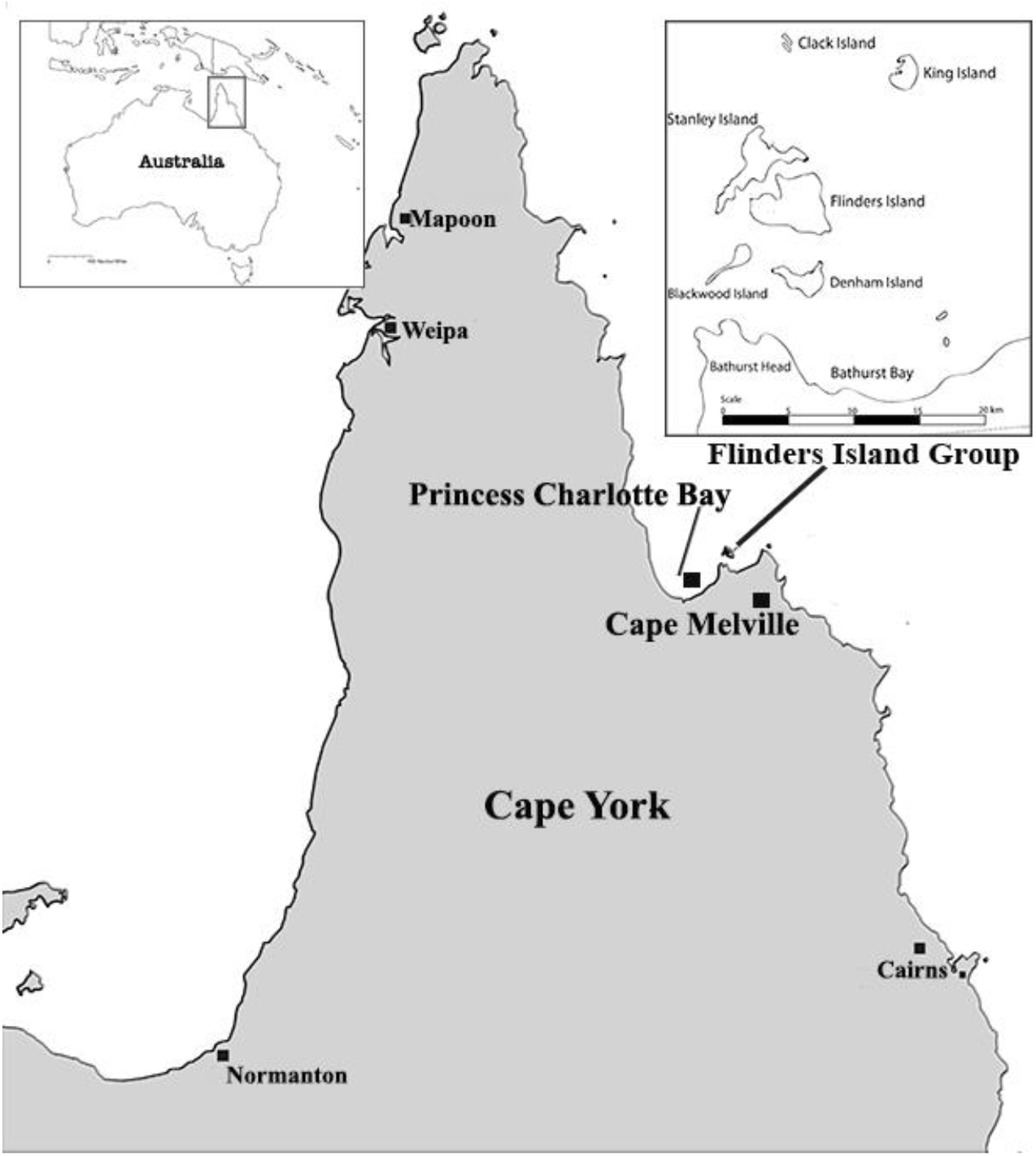
Map of Cape York, showing. The Flinders Island group location in the tropical north-east of Queensland off the eastern coast of Cape York.

Princess Charlotte Bay was first recorded by British navigators in 1815 (Jack, 1921). Princess Charlotte Bay was situated within the early shipping routes of the east coast of Australia, allowing early contact between Europeans and Traditional Owners in the 19th Century (Coppinger, 1883; King, 1827; Roth, 1898). Although there has been extensive archaeological research conducted in the Flinders Group, no research to date has extended to mortuary practices. Little attention has been paid to many of the burial sites on the islands, or indeed throughout many parts of Cape York. Archaeological examinations carried out by Beaton in 1985 revealed that there has been an Aboriginal occupation of the Flinders group for at least 2,300 years (Beaton, 1985). Beaton also suggested that the initial occupation of Flinders Island occurred ~2500 years ago and was probably closely related to the introduction of Papuan/Melanesian outrigger canoes (Beaton, 1985). This interpretation is now under question with the discovery of archaeological evidence at Endean Shelter illustrating that Aboriginal occupation of the islands extended back to 6,280 calBP (Collard et al., 2019). Ethnographic records provide detailed information on mortuary protocols after initial contact, but little is known about how these customs may have differed in pre-European contact.

Knowledge of Aboriginal mortuary practices in north-east Australia is limited to ethnographic accounts by anthropologists and observers in the 19th and early 20th Centuries (Roth, 1898, 1907). According to Roth, people of Cape York interpreted death as a result of spiritual intervention or human agency, rather than natural phenomena. It was believed that spirits of the dead could harm the living (Roth, 1907). The deaths of prominent and/or powerful people were often avenged by their remains being carried from camp to camp and defleshed before finally being buried or interred in trees or caves. While old, less esteemed, or infirmed people were given simpler burials with minimal ceremony and often buried within close proximity to the site of death (Roth, 1907). Also, at Torilla, south of Princess Charlotte Bay, Roth observed that women were usually buried immediately after death, bundled in bark and carried from camp to camp (Roth, 1907). In this study, we propose a mortuary narrative constructed from genomic data recovered from five ancient individuals who were interred in the Flinders Island Group.

We suggest that our understanding of the mortuary practices of Aboriginal Australians can be improved using ancient DNA methods. We investigate the possible link between mortuary practices and kinship among five individuals excavated within the Flinders Island Group in 2015-2016 (Adams et al., Submitted). Three burials were located within rock-shelters and two on beaches. The two contrasting sites pose an interesting question about the kinship among individuals, and at different sites. To gain a direct insight into the kinship relationships we used complete mitochondrial genome sequences and Y chromosome data, whenever it was available. Although we obtained uniparental genomic data by applying whole-genome target enrichment coupled with Next Generation Sequencing (NGS) methods, the autosomal data is not the focus of this publication.

## 2 Materials and Methods

### 1.1 Archaeological samples

Archaeological fieldwork included rescue excavations of two disturbed beach burials and recording of three bundle burials from two interment caves in the Flinders Islands (Adams et al., Submitted). The research focused on investigating burial sites and recovering samples for paleogenetic and isotope research. Orientation, shape, and size of each burial were recorded. Anatomical measurements were used for preliminary determination of sex, age, ancestry, and pathologies (Adams et al., Submitted). All observation and recording of available skeletal elements were completed in the field following excavation (Adams et al., Submitted). Immediately upon completion of recording and sampling Traditional Owners reinterred all excavated elements at a safe location. With consent from Traditional Owners, tooth and bone samples were collected for radiometric dating, isotopic assessment and aDNA analyses.

#### 1.1.1 Details of the individuals studied

##### Flinders Island individual 1 (FLI1)

this adult male was discovered eroding from a beach burial in an area of Flinders Island known as Apa Spit (Hale and Tindale (1933) or Wathirrmana (Sutton et al., 1993). The remains were initially excavated by Traditional Owners and Queensland Police, who determine that it was a traditional burial, at a depth of ca. 1.2 m below the original ground level (Adams et al., Submitted). The individual has a north-east orientation with their face directed east. They had been interred on their back with their legs partially flexed and a large (ca. 40 cm diameter) limestone rock placed on his chest. The individual’s hands were placed palm down on the thighs, and their feet were crossed. The rock, which was removed during the preliminary investigation, was the only identifiable grave good (Adams et al., Submitted).

##### Stanley Island individual (STI1)

In 2015, this female was discovered eroding from the beach foredune sands on Stanley Island by two fishermen, who removed the crania for a photo before re-burying the remains (Adams et al., Submitted). The burial was located in a large (ca. 1 km wide), flat, sandy cove that is surrounded by limestone boulders. The individual had been buried facing south-east. Wear of the frontal and supraorbital ridges of the cranium indicating a long-time exposure of the remains.

##### Flinders Island individual 2 (FLI2)

a set of remains found inside a rock-shelter located close to the beach on the east coast of Flinders Island. The rock-shelter faces east and is ca. 10 m wide and ca. 2 m deep. Graham Walsh, in his early survey of the islands in the 1980s, recorded two sets of remains in the rock-shelter (Walsh, 1985) suggesting that they were bundle burials. Only one set of these was found belonging to an adult male (FLI2), to be still existing in the rock-shelter which was partially covered as a result of heavy weathering of the roof.

##### Flinders Island individuals (FLI3 (B2) and FLI4 (B3))

the second rock-shelter is located on the north coast of Flinders Island. It faces north-west and is ca. 30m long, ca. 6m deep, and up to 1.5 m high. Again, the rock-shelter was surveyed by Walsh who recorded the presence of six bundle burials involving ornate bark coffins and matting. In 2016, only two of the burials remained Walsh’s (Walsh, 1985) Burial 2 (B2) of a young male and Burial 3 (B3) of an early 20’s female (**Figure 2D and 2E**). Although the cylinder-coffins and skeletal remains associated with four of the six burials were missing, the outer paperbark wrapped around the cylinders remained either *in situ* or nearby, except for Burial 6 which had no evidence of discarded paperbark. One cylinder-coffin remained with FLI4 (B3). It had been opened but retained fine twine that likely bound the post-cranial skeletal elements.

**Figure 2.**
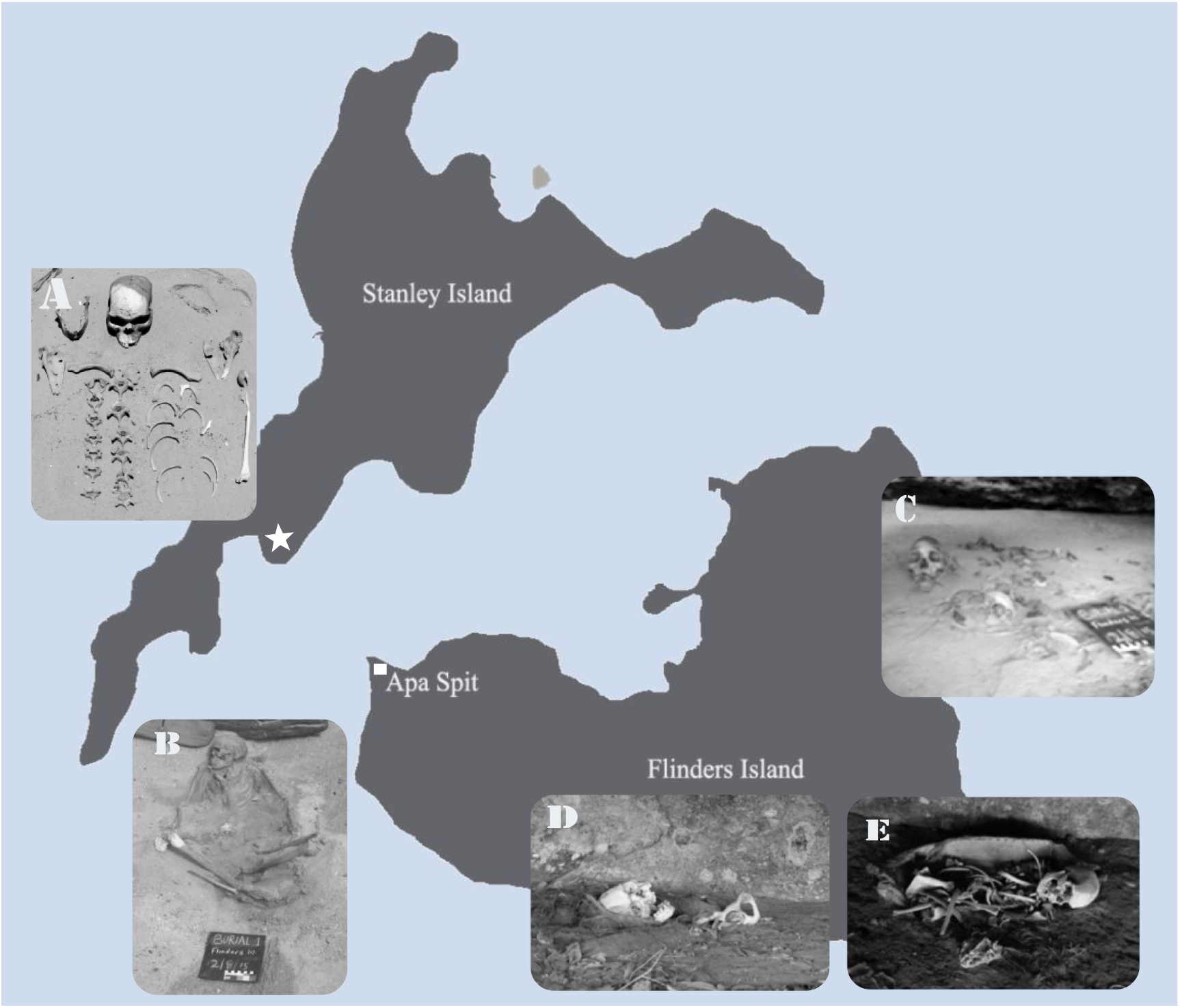
Flinders Island Group burials. **2A)** Stanley Island female (STI1) remains, the burial location is indicated by the hexagon on the map. **2B)** Flinders Island adult male individual (FLI1) beach burial in the area of Apa Spit. **2C)** A set of remains found inside a rock-shelter (FLI2), located close to the beach on the east coast of Flinders Island. The second rock-shelter in the north coast of Flinders island where **2D)** FLI3 and **2E)** FLI4 both were buried.

### 2.1 Radiocarbon dating

Human bone collagen from each of the five sets of remains was directly dated using AMS radiocarbon methods. This procedure was conducted at the Research School of Earth Sciences, Australian National University, Canberra. Dates were calibrated using OxCal 4.2 (Bronk Ramsey, 2013) software and the Southern hemisphere calibration curve SHCal13 (Hogg et al., 2013) (**Table S3).**

### 2.2 Ancient DNA

Ancient DNA work was carried out in a dedicated facility in the Australian Research Centre for Human Evolution at Griffith University. Only the roots of teeth were used for the ancient DNA extraction of the five individuals. Methods for handling ancient DNA were as outlined by Knapp et al.(Knapp et al., 2012).

Each sample was initially decontaminated with 10% sodium hypochlorite for 10 min, followed by 80% ethanol, and five minutes under UV light. Subsequently, the skeletal material was processed using a Dremel rotary tool with a high-speed diamond cutter head, or manually with a sterilised scalpel blade. DNA was extracted from ~50mg of bone or tooth powder following the modified protocol outlined in Wright et al. (Wright et al., 2018), which allowed for better recovery of shorter DNA fragments (~30 bp). Negative controls were included throughout all procedures, each of which showed no contamination.

#### 1.1.2 DNA library construction methods

Double-stranded Illumina DNA libraries were built according to the modified method of Meyer and Kircher (Meyer & Kircher, 2010) as detailed in Wright et al. (Wright et al., 2018). Using the NEBNext DNA Library Prep Master Mix Set for 454 (New England Biolabs ref: E6070) 21.25μl of DNA extract was subjected to three consecutive steps: NEBNext end repair, NEBNext blunt end adaptor ligation, followed by an Adapter Fill-In reaction. A MinElute (Qiagen) purification step with 10x binding buffer PB (Qiagen) was carried out between the first and second steps.

All pre-PCR libraries were amplified using KAPA HiFi Hotstart Uracil+ (Kapa Biosystems), according to manufacturer’s instructions using Illumina single indexing primers. PCR amplification products were cleaned using 1x Axygen beads according to the manufacturer’s instructions. Amplified libraries, including negative controls, were quantified and visualised for length distribution using the 5000 High-Sensitivity DNA tapes on the TapeStation 4000 (Agilent Technologies), following the manufacturer’s instructions.

#### 1.1.3 Whole-genome in-solution target capture

Between 100 – 500ng of library amplified DNA was subjected to in-solution target enrichment using whole human genome myBaits^®^ WGE (Arbor Biosciences) as detailed in Wasef et al. (Wasef et al., 2018) and Wright et al. (Wasef et al., 2018; Wright et al., 2018). Target capture enrichment steps were performed according to manufacturer’s instructions with the following modifications: the hybridisation step was performed for 36 – 42 hrs at 57°C. The beads, and the bead binding buffers were heated to 57°C for 30 min before being used. Further cleaning steps were also performed at the same hybridisation temperature. Post-capture libraries were amplified on beads using HiFi HotStart Uracil+ ReadyMix (Kapa Biosystems) for between 14 and 17 cycles, and then visualised using the 5000 High-Sensitivity DNA tapes on the TapeStation 4000 (Agilent Technologies).

#### 1.1.4 Ancient sequencing

After target enrichment, ancient samples were sequenced on HiSeq 4000 Sequencing System (Illumina) at The Danish National High-Throughput DNA Sequencing Centre in Copenhagen. Sequences were base called using CASAVA 1.8.2 (Illumina), demultiplexed and FASTQ files were generated by the sequencing facility.

### 2.3 Modern Aboriginal genomes

In addition to the previously published mitochondrial genomes of Indigenous Australians (Hudjashov et al., 2007; Nagle et al., 2017; Rasmussen et al., 2011; Tobler et al., 2017; van Holst Pellekaan et al., 2006), we also incorporated the haplogroup data previously published in (Malaspinas et al., 2016; Wright et al., 2018) (**Table S2**).

### 2.4 Genome analyses

Adapter sequences were trimmed using fastx_clipper, part of FASTX-Toolkit 0.0.13 (http://hannonlab.cshl.edu/fastx_toolkit/), with reads shorter than 30 bases and low-quality bases removed using parameters -Q 33 −1 30. Reads were aligned to the human reference build GRCh37/hg19 for the nuclear genome or to the revised Cambridge Reference Sequence (rCRS) (accession number NC_012920.1). For mitochondrial data BWA 0.6.2-r126 software was used to align sequences (R. Li et al., 2010) with the following options: seed disabled (Schubert et al., 2012) and terminal low-quality trimming (using parameter -q15). Duplicate reads were removed using the MarkDuplicates tool from the Picard 1.68 tools package (http://broadinstitute.github.io/picard/). The mapped reads were sorted, indexed and merged using SAMtools 0.1.18 (H. Li & Durbin, 2009, 2011). The consensus mitogenome was generated using the SAMtools bcftools view –cg -command and converted to FASTA via SAMtools/bcftools/vcfutils (H. Li & Durbin, 2009). Qualimap was used to estimate the levels of coverage (Okonechnikov et al., 2015) and the number of mapped reads to the human reference genome (GRCh37/hg19).

Ancient DNA sequences were authenticated using MapDamage software (Jonsson et al., 2013), which uses levels of cytosine to thymine misincorporations in the 5’ end of fragments, and guanine to adenine misincorporations in the 3’ end. Schmutzi software was used to estimate modern human contamination levels in the ancient mitochondrial sequences using the contDeam command which estimates contamination levels using deamination patterns (Renaud et al., 2015). Endogenous consensus sequences were generated using default settings. Both the Schmutzi generated consensus sequences and the original ancient sequences were then manually checked using the SAMtools tview command (H. Li & Durbin, 2009). Missing sites were replaced with “N”. ANGSD was also used to estimate modern contamination levels in male samples.

Mitochondrial haplotypes were identified using HaploGrep 2.0. A total of 229 ancient and modern mitochondrial genomes (as detailed in Table S2) were realigned using the online version of MAFFT software (Katoh et al., 2017). The mitochondrial consensus sequences were used to construct a Maximum Likelihood phylogenetic tree using the online version of RAXML with 1000 bootstrap replications (Kozlov et al., 2019). The resulting Likelihood tree provided information about the maternal ancestry of each of the Flinders Island Group individuals (Figure 3).

**Figure 3.**
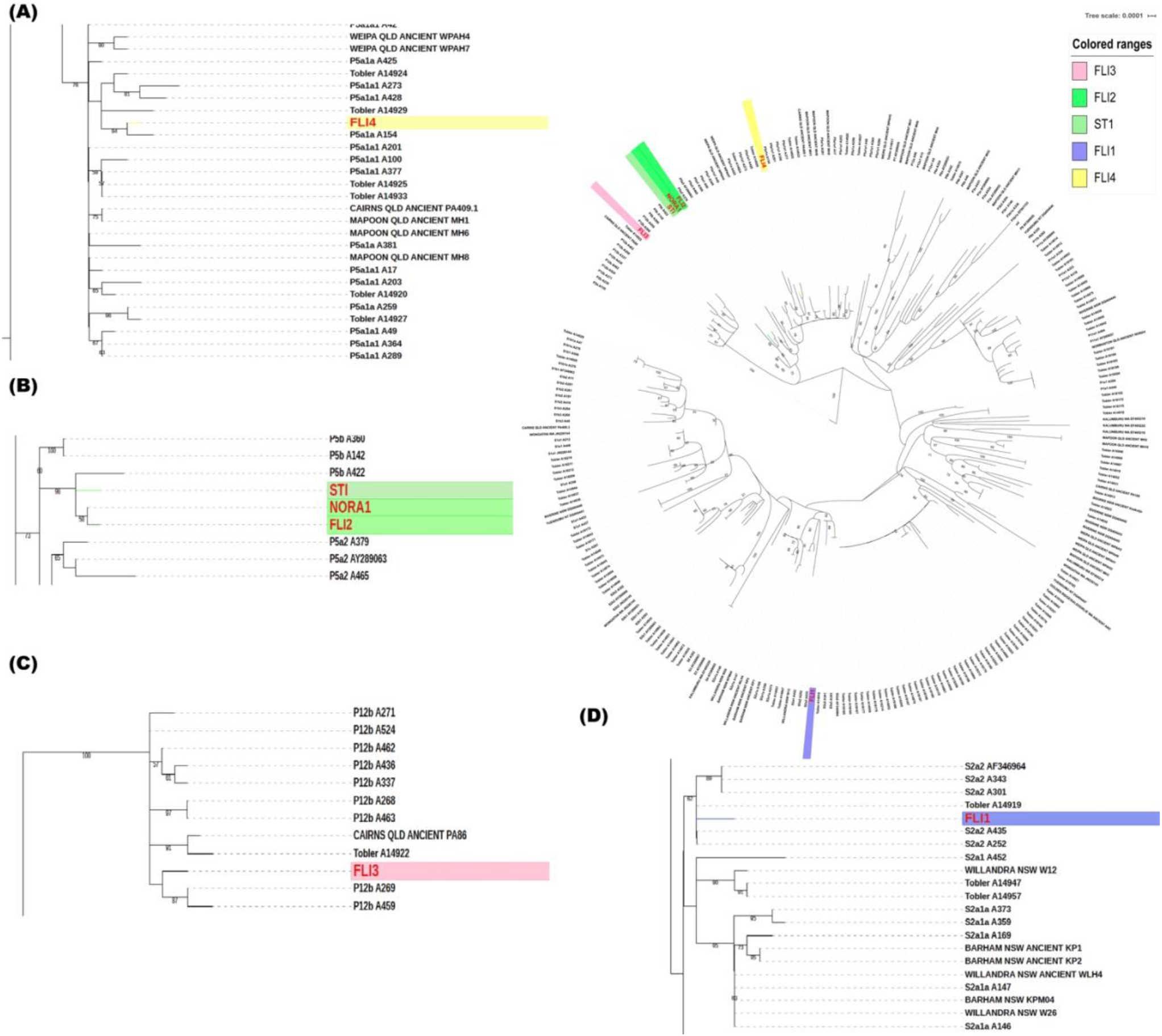
Mitochondrial maximum likelihood phylogenetic tree. **A)** FLI4 belongs to the P5a1a haplotype. **B)** FLI2 and STI1 both belong to mitochondrial haplotype P5b1. **C)** FLI3 belongs to the P12b haplotype. **D)** The FLI1 showed a S2a haplotype.

Summary statistics of haplogroup frequencies in Queensland were estimated using 144 mitogenomes summarised in Table S4. Arlequin software V3.5.2.2 (Excoffier & Lischer) was used to estimate haplotype frequencies and genetic distances (Fst) as pair-wise values, and to perform analysis of molecular variance by means of AMOVA (Table S4 & S5). Using the SPSS V26.0.0.1 software package, Principal Component Analysis (PCA) was performed on the haplogroup frequencies detected in the Queensland populations investigated and in previously studied populations (Figure S1A). Fst distance matrices of mtDNA haplotypes were used to construct MDS plots (Figure S1B). Median-joining networks of haplogroups without pre- and post-processing steps were performed using Network (http://www.fluxus-engineering.com/) (Figure S2).

Sex determination of all ancient Aboriginal Australian individuals was inferred using the method outlined in Skoglund et al. (Skoglund et al., 2013). Y chromosome haplogroup assignments were performed for male individuals using Yleaf software (Ralf et al., 2018).

## 3 RESULTS

### 3.1 Radiocarbon dating

Supplementary Tables S1 and S3 include the AMS 14C dates conducted on the bone collagen of each of the five individuals. The remains were dated between 147 and 473 calBP (**Table S3**). These results show that all individuals recorded on both Flinders and Stanley Islands died before European colonisation, making them suitable for comparison with the ethnographic mortuary record.

### 3.2 Ancient DNA

We successfully recovered complete ancient mitogenomes from five ancient Flinders Group individuals, in addition to 29 we published previously (Wright et al., 2018), ranging between 2.3 and 331.9x coverage (**Table S2**). All ancient DNA recovered were authenticated, and modern-day contamination levels were estimated (**Table S1**). The characteristic aDNA damage patterns were estimated for each sample using MapDamage software (Jonsson et al., 2013). All samples exhibited damage patterns characteristic of ancient DNA, with elevated levels of cytosine to thymine misincorporations in the 5’ end of fragments, and guanine to adenine misincorporations in the 3’ end (Dabney et al., 2013). The mean read length also indicated that the DNA recovered was likely authentic (**Table S1**). Recovered sequences were also consistent with Aboriginal Australian mitochondrial haplotypes and did not match those carried by any of the ancient DNA laboratory members.

Sex determination of the five individuals was determined bioinformatically, using the method detailed in Skoglund et al. (Skoglund et al., 2013) and confirmed that FLI1, FLI2, FLI4 were males, while STI1 and FLI3 were females.

### 3.3 Genome analyses

After constructing a Maximum Likelihood phylogenetic tree using 229 ancient, historical and modern mitochondrial genomes (**Table S1, S2 and Figure 3**), it became clear that the five ancient individuals fell within previously described mitochondrial haplotypes of contemporary and ancient Aboriginal Australians (Hudjashov et al., 2007; Malaspinas et al., 2016; Rasmussen et al., 2011; Tobler et al., 2017; van Holst Pellekaan et al., 2006; Wright et al., 2018). All five mitochondrial genomes were Aboriginal Australians in origin. In the network generated, FLI1 clustered with two individuals from Central and South Queensland, both carrying the S* haplogroup (Supplementary Figure S2)

112 unique haplotypes were present among the 124 mitochondrial genomes from Queensland, showing high haplotype diversity (Hd= 0.9979). When comparing the haplotypes within Queensland, the Flinders Island group ancient samples showed a high mtDNA diversity (Hd= 1.000). Analysis of molecular variance based on haplogroup frequencies demonstrating the variation among different QLD populations are summarized in Table S5.

FLI2 and STI1 have the recently identified Aboriginal mitochondrial haplotype P5b1 (Wright et al., 2018), which is between 12,000 and 28,000 years in age and appears to be restricted to Australia. Ancient Aboriginal Australians FLI2, STI1, NORA1 from Normanton, and the previously published A422, a contemporary individual from Queensland, all carry the P5b1 mitochondrial haplotype (**Figure 3A**). These four mitogenomes are the only Aboriginal Australians included in this research that carry the P5b1 haplotype. Ancient individual FLI4, found in the same rock-shelter as FLI3, carries the P5a1a haplotype, which is also present in contemporary Aboriginal Australians from Queensland (**Figure 3A**). FLI3 carries the mitochondrial haplotype P12b, also carried by 20 other individuals from Queensland. P12b representing the highest observed haplotype in Queensland with 13.4% (Table S4 and Figure S3)

Unexpectedly, ancient individual FLI1 carries a S2a mitochondrial haplotype. This haplotype indicates a maternal ancestor for that individual who was not from the Flinders Island group or any close mainland community, but rather this haplotype is more closely related to haplotypes found in New South Wales, central Queensland and South Australia (Table S4 and S5, **Figure 3C and Figure S3**).

### 3.4 Y-chromosome

Few Y-chromosome studies of Aboriginal Australians have been published to date, with the majority showing unique Aboriginal Australian Y-chromosome haplogroups, C* and K* predominantly, and M* in rare cases. As a result, there is a limited modern Y-chromosome database, which did not allow for phylogenetic analyses similar to the work done on the mitochondrial genomes (Bergstrom et al., 2016; Nagle et al., 2017). In previous studies, one constant, however, was the detection of considerable levels of Eurasian admixture in modern individuals, with a large number of research participants self-identifying as Aboriginal Australian carrying non-Indigenous Y-chromosome haplotypes. The level of Eurasian admixture varied from study to study, with ~32% being reported by Malaspinas, et al. (Malaspinas et al., 2016), ~56% by Nagle, et al. (Nagle et al., 2016) and ~70% by Taylor et al. (Taylor & Henry, 2012).

FLI2 showed the S1c haplotype (characterised by Z41926, Z41927, Z41928, Z41929, Z41930 SNPs), while FLI4 belongs to the S1a3a haplotype (which is a subclade of S1a~ previously known as K2b1a). Both of these haplotypes are unique to Aboriginal Australians. The determination of FLI1’s haplotype was not possible due to the low coverage of the Y chromosome.

## 4 DISCUSSION

The ethnography of the Flinders Islands as discussed by Hale and Tindale (Hale & Tindale, 1933) suggests that the status of an individual in life was reflected in the complexity associated with their mortuary practices. Moreover, the Flinders Island group were often visited by people from the mainland and other islands. During their visits to the island they were often involved in ceremonies (Hale & Tindale, 1933; Rigsby & Chase, 1998). A map of Apa spit, where individual FLI1 was found eroding from the beach, drawn by Tindale during his visit to Flinders Island in the 1920s, showed visitation to the islands by at least four other tribal groups at that time. Visitors and non-locals were also offered different mortuary practices.

FLI1 was an older individual exhibiting extreme occlusal wear and periapical lesions (Adams et al., Submitted). Being an elderly individual at the time of death FLI1 may have represented one of the less esteemed members of the community that was not seen as a threat in the afterlife. He was buried a short time after death with no signs of extensive ceremony and complex interment. The rock placed on the torso of FLI1 represents a distinct funerary practice hitherto undescribed in available published literature for Australia. No other grave goods were observed in this burial. A modern Aboriginal interpretation of the purpose of the stone as a grave object, was provided by Traditional Owner Danny Gordon, who commented that the beach burial of FLI1 may have been an ‘unliked man’(Danny Gordon pers. comm 2015). Alternatively, the FLI1 man may have died during his visit to the island and hence represents an example of a beach burial afforded to non-locals, as recorded by Hale and Tindale (Hale & Tindale, 1933). The mitochondrial genome of FLI1 provided more insights into his possible ancestry. This individual carries a mitochondrial haplotype (S2a) that is more dominant in New South Wales, especially among the Willandra Lakes communities (**Table S2, S4 and Figure S3**). We showed that the male beach burial (FLI1) carried a mitochondrial lineage that differed from other individuals, including contemporary communities from North Queensland. The haplotype difference of FLI1 from the other Flinders Island individuals may suggest that he was born away from the island but raised there, or was a visitor to the island at the time of his death.

The young woman (STI1) was also recovered from beach foredunes, so she might have been another example of a visitor burial. However, the recorded correlation between social status and the extent of ceremony and burial complexity is perhaps another explanation for the nature of her internment. Her placement on the beach indicating that this young woman did not meet the criteria for a more complex burial ritual (Adams et al., Submitted; Roth, 1907). She was buried articulated, and therefore not defleshed before burial, with no other burial goods found. However, it is important to note that this burial had been heavily disturbed, so it is possible that such goods may have existed prior to the disturbance of the burial (Adams et al., Submitted). Notably, we observed that STI1 carries a maternal haplogroup (P*) that dominates Aboriginal Australian communities in Queensland (**Figure S3**), especially Cape York. particularly Cape York. Moreover, the STI1 woman shared an ancestral maternal lineage with the FLI2 individual who was buried in the rock-shelter. These results strongly suggest that the STI1 woman was likely a Flinders Island group individual, however, we could not rule out the possibility that she was a visitor from the nearby mainland.

Bark burial coffins are a feature of the Flinders Islands burial landscape, indicating a more complex form of funerary practice. All three sets of remains (FLI2, FLI3 and FLI4) recorded here represent individuals that were ritually prepared in accordance with burial protocols that have been ethnographically documented (Hale & Tindale, 1933; Roth, 1898, 1907). The genomic data of those three Flinders Island Aboriginal Australians, combined with modern genomes, form the basis of our understanding of the maternal haplotypes expected from that area of Queensland. Although the three belong to a common haplogroup (P*) for the region, they are from three different sub-haplotypes; P5b1, P12b and P5a1a consecutively.

The absence of a close kinship relationship between FLI2 and FLI4 was supported by the little information gained through the study of the paternal lineage. Both FLI2 and FLI4 carried two different Indigenous Australians Y-chromosome haplotypes (S1c and S1a~). Moreover, the C14 dates for those two individuals do not overlap after the Marine13 correction (Table S1). This reveals that FLI3 (B2) and FLI4 (B3), despite being buried within the same cave, did not share a kinship relationship.

However, the Y chromosome results, when compared with the contemporary population from Queensland, did not provide any additional understanding of the burial practices performed on the island. This was a result of the significant levels of Eurasian admixture observed in the contemporary population.

## 5 CONCLUSIONS

Although burials of Indigenous people often represent a poor source of DNA preservation, particularly in Australia, we have shown here that it is possible to recover full mitochondrial genomes, even in tropical contexts. The analysis of the ancient mitogenomes, in combination with a fine-scale genomic map of Aboriginal Australian mitochondrial haplotypes, can be used to better understand and test a range of hypotheses about mortuary practices and changes over time. Our genomic findings resolved to provide indicative answers to the questions about the beach burials, showing that FLI1 who was buried on the beach, most likely, for being a non-local, rather than being a less esteemed member of the Flinders Island community. While the STI1 female was more likely a less esteemed member of the community. These results do reflect some of the purported inequalities in Indigenous Australian mortuary practices that might have been based on age, sex or status of individuals, but they also reveal that ethnohistoric observations may reflect biases from early ethnographers. The presence of a young female FLI3 buried in the rock-shelter, indicates that both sexes were afforded complex mortuary rituals by their community.

## 8 Author Contributions

SW, MW and DL conceived the project. SW, CF and MW conceived the idea of the manuscript. DL, MW and EW provided funding. MW, SA and CF conducted the field work and collected ancient samples from the Flinders Island Group. SW performed the experiments and analyses of the data. JW and SW checked the mitochondrial haplotypes. SW interpreted the data with help from DL, MW, CF and JW. SW drafted the manuscript. JW, DL and SW revised the paper, with contributions from all the authors.

## 10 Acknowledgments

We thank all the Aboriginal Australian participants for supporting this research. We would like to acknowledge the QLD Police, DATSIP and QPWS for assistance in the field. We also thank David McGahan for his assistant in the excavation of Stanley Island. We thank the National Throughput DNA Sequencing Centre at the University of Copenhagen for ancient DNA sequencing.

## 11 Supplementary Material

The Supplementary Tables S1-S5 and Figure S1-S3 for this article can be found online at:

## 12 Data Availability Statement

Ancient Aboriginal mitochondrial data have been deposited in the GenBank database (http://ncbi.nlm.nih.gov/genbank), which is hosted by the National Center for Biotechnology Information, under accession numbers MK165665 to MK165690 (more specifically MK165683.1, MK165666.1). All data needed to evaluate the conclusions in the paper are present in the paper and/or the Supplementary Materials.

